# Disentangling Conditional Effects of Multiple Regime Shifts on Atlantic Cod Productivity

**DOI:** 10.1101/2020.07.28.224477

**Authors:** Tommi Perälä, Esben M. Olsen, Jeffrey A. Hutchings

## Abstract

Regime shifts are increasingly prevalent in the ecological literature. However, definitions vary, and many detection methods are subjective. Here, we employ an operationally objective means of identifying regime shifts, using a Bayesian online change-point detection algorithm able to simultaneously identify shifts in the mean and(or) variance of time series data. We detected multiple regime shifts in long-term (59-154 years) patterns of coastal Norwegian Atlantic cod (>70% decline) and putative drivers of cod productivity: North Atlantic Oscillation (NAO); sea-surface temperature; zooplankton abundance; fishing mortality (*F*). The consequences of an environmental or climate-related regime shift on cod productivity are accentuated when regime shifts coincide, fishing mortality is high, and populations are small. The analyses suggest that increasing *F* increasingly sensitized cod in the mid 1970s and late 1990s to regime shifts in NAO, zooplankton abundance, and water temperature. Our work underscores the necessity of accounting for human-induced mortality in regime shift analyses of marine ecosystems.

## Introduction

The productivity of commercially valuable marine species is a consequence of the direct and interactive effects of human-induced mortality (exploitation, habitat destruction), natural environmental shifts, and climate change [1]. Fundamental to our understanding of these synergistic processes is the concept of regime shifts [2]. A common approach is to describe, through one means or another, an abrupt temporal change in a measure of biological productivity as a regime shift and to then explore other data to identify causal drivers of the shift. Temporally abrupt changes in biological productivity are clearly of importance from basic ecological and management perspectives, as they can affect population dynamics, species viability, ecosystem structure and function, and fisheries sustainability [3–4].

However, what constitutes a regime shift is not always clear. Many definitions explicitly refer to ecosystems and incorporate the necessity that a shift from one regime to another must be difficult to reverse, asserting that regimes represent stable alternative states in community structure [5] or ecosystem configuration [6].

A second issue is methodological in nature. Some analyses are based on empirical but ultimately subjective impressions resulting from visual observation, such as an over-grazed kelp bed [7] or coral bleaching [8]. When evidence of a regime shift is visually striking and arguably self-evident, there is perhaps less need for the methodological objectivity that long-term, continuously variable information usually demands. Under such circumstances, sequential *t*-tests are not uncommon [9–12], although the selection of years applied is frequently based on the researcher’s perception of the magnitude of the effect of the purported regime shift.

We offer an alternative, operationally objective means of identifying regime shifts in time-series data that involves application of a Bayesian online change point detection (BOCPD) algorithm [13]. Perälä and Kuparinen introduced this approach to detect regime shifts in the fisheries ecology literature [14]. Their method utilized normal-gamma conjugate priors for the normal observation model, resulting in an analytically tractable procedure that is able to simultaneously detect shifts in the mean and(or) variance parameter of the data-generating process. Using sequential Monte Carlo (SMC) methods, Perälä et al. expanded the methodology so that it can be used to detect shifts in any parameter of the underlying predictive model with arbitrary prior distributions [15]. They applied the method to Beverton-Holt and Ricker fisheries stock-recruitment models and detected shifts, for example, in maximum per capita reproductive output parameters.

The BOCPD algorithm continually and sequentially updates (i) the posterior probability distribution since the latest regime shift together with (ii) the posterior probability distributions of the parameters of the data-generating process. A high probability of a change point (regime shift) results from poor compatibility of the model prediction with an observed data point, as determined by the posterior predictive density function evaluated at the new observation. Here, we use the SMC implementation of the algorithm for a normal observation model with uniform priors for the mean and variance parameters of the model.

Our overarching objective is to use the BOCPD algorithm to detect regime shifts in a measure of biological productivity (a nearly century-long time series of juvenile Atlantic cod, *Gadus morhua,* abundance) and hypothesized drivers of cod productivity [3, 16–19]: North Atlantic Oscillation (NAO); zooplankton abundance; and water temperature. The NAO reflects changes in environmental factors that have direct effects on productivity, such as wind strength, currents speeds, temperature, and turbulence [20]. We use the BOCPD algorithm to identify regime shifts in all time series. The degree to which these time series overlap with one another will be used to infer potential causal drivers of regime shifts in cod productivity. In addition to these metrics of climate, food supply, and temperature, we applied the BOCPD algorithm to temporal estimates of fishing mortality to explore its role as a driver [21] and as a factor that might conditionally affect the strength of other suspected drivers.

## Materials and methods

The BOCPD algorithm [13] is based on sequential Bayesian posterior estimation of the length of the current regime (run length) and the regime-specific parameters of the underlying predictive model (mean and variance). The recursion for calculating the posterior distribution of the run length *r*_*t*_ at time *t* given the observations *y*_1:*t*_ = (*y*_1_,…,*y*_*t*_) is

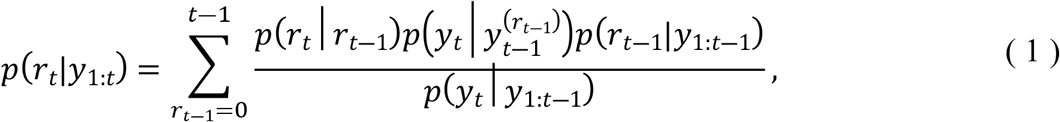

where *p*(*r*_*t*_|*r*_*t*−1_) is the change point prior distribution containing our prior beliefs about the probability of a regime shift. We assume constant prior probability for a shift defined by the hazard rate parameter *λ*,

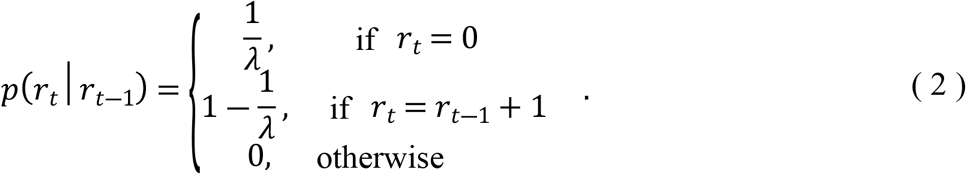

The underlying predictive model (UPM), 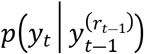, is the predictive density conditional on the run length and thus on the latest *r*_*t*−1_ observations, i.e., 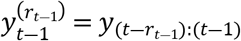. The UPM is used to give more weight to those run lengths that better predict the new observation, and it is defined by the observation model,

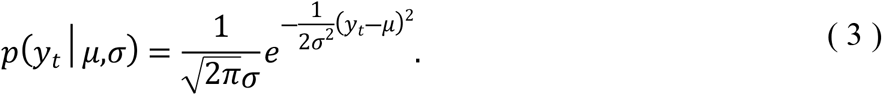

If none of the existing runs explains the new observation well enough, the prior predictive density, 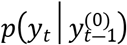, will dominate, resulting in a high posterior probability for a regime shift.

As we are mostly interested in retrospective analysis of regime shifts, we use the smoothed run length probabilities to find the most likely segmentation of the data or the most likely set of regimes by maximizing the product of run length probabilities over the whole time series [15]. We can find the maximum among all possible combinations of regimes or we can focus on some subset of regimes by setting certain constraints for the regimes. We have decided to set a constraint for the minimum length of the regimes (*M*). This constraint is not used for the first and the last regimes, though, since their start and end points can be outside the time frame of our data. We use uniform priors for the mean and variance parameters in each of the time series analyzed. The lower and upper limits for the uniform priors were assigned so that all plausible parameter values were contained in the intervals. The posterior inference of the observation model parameters is carried out by a sequential Monte Carlo algorithm [15], using 100,000 particles.

### Model parameters

There are two key parameters in the regime shift detection method. The first is the minimum length of each regime, *M* (*M*=10 would imply a minimum regime length of 10 year). For some data sets, *M* can be informed by prior knowledge. The suggestion has been made, for example, that the NAO data exhibit decadal variability (e.g., [22]). This might provide a defensible ‘default’ value of *M*=10.

In general, we can see a cogent argument for maintaining *M* at 10 years for all data sets primarily to ensure that we are not biasing analyses against the detection of significant, but comparatively brief, regimes. In addition to conforming with general patterns in the NAO [22], ten years approximates one generation for northeast Atlantic cod (in the absence of fishing), and should be sufficiently long to detect persistent shifts in copepod abundance and water temperature.

The second parameter of import is the hazard rate in the change point prior, λ. Lambda can be thought of as the probability or expectation of the frequency of a regime shift, e.g., roughly every 10 years (λ=10) or every 25 years (λ=25). This parameter is more challenging to base on empirical expectation. Nonetheless, some caution in setting its value can be exercised. Relatively small values of λ (e.g., ‘10’) might lead one to falsely interpret short-term periods of seemingly extreme, potentially spurious values as regimes. On the other hand, relatively large values such as λ=50 (regime shifts occurring twice in each century) might be considered unduly ‘conservative’, biasing against the detection of biologically meaningful regime shifts because of the use of a prior that constrains regime shifts to be comparatively infrequent.

For our analyses, we set *M*=10 for all data sets. For the hazard rate, we compare model outputs for λ=10, 20, 25, and 50. Outputs for all variables at alternative values of λ are presented in the Supplementary Information.

### Data

Our measure of biological productivity reflects the abundance of juvenile Atlantic cod (ages 0+ to 2+ yr) in Skagerrak, a strait running between the southeast coast of Norway, the west coast of Sweden, and the Jutland peninsula of Denmark. Standardized beach-seine surveys have been conducted along coastal Norwegian Skagerrak annually since 1919 [23]. The time series of data available for our analyses extended from 1919 to 2014.

In addition to cod, we examined time-series data for hypothesized drivers of cod productivity. Winter NAO data (December through March) were obtained for the years 1864 through 2018 [24]. As a measure of juvenile cod food supply, we tested for the presence of regime shifts in the abundance of the energy-rich calanoid copepod *Calanus finmarchicus,* estimated by the Continuous Plankton Recorder (CPR) survey for the survey area closest to, and overlapping with, the Norwegian Skagerrak (area C1) (https://www.cprsurvey.org/data/data-charts/) [25]. We pooled the CPR data for the months March through August, the period during which zooplankton are available to, and consumed by, cod larvae and juveniles in the North Sea [26]. Sea-surface water temperatures in Skagerrak are available from the Flødevigen Research Station, Institute of Marine Research (http://www.imr.no/forskning/forskningsdata/temperatur_flodevigen/draw.map?boey=1). Temperatures recorded at Flødevigen are positively correlated with temperatures elsewhere in Skagerrak (e.g., [16]). These data were analysed on a monthly basis for the years 1925 to 2017. Estimates of fishing mortality (*F*) and spawning stock biomass (SSB) are available for North Sea cod, the management unit of which Skagerrak cod is a part [27].

## Results

### North Atlantic Oscillation

The BOCPD algorithm (at λ=10 and *M*=10) produces a pseudo-decadal pattern in the winter NAO index, indicating that the model is capturing the temporal dynamics previously ascribed to this index [22] (Fig 1a). Over the 140-year time series, ten regimes are identified. Some, such as those between 1916 and 1960, are comparatively brief, raising questions as to whether these patterns in the mean and(or) variance in the data are consistent with some stipulations that regime shifts represent stable states that are difficult to change [5–6].

**Fig 1.**
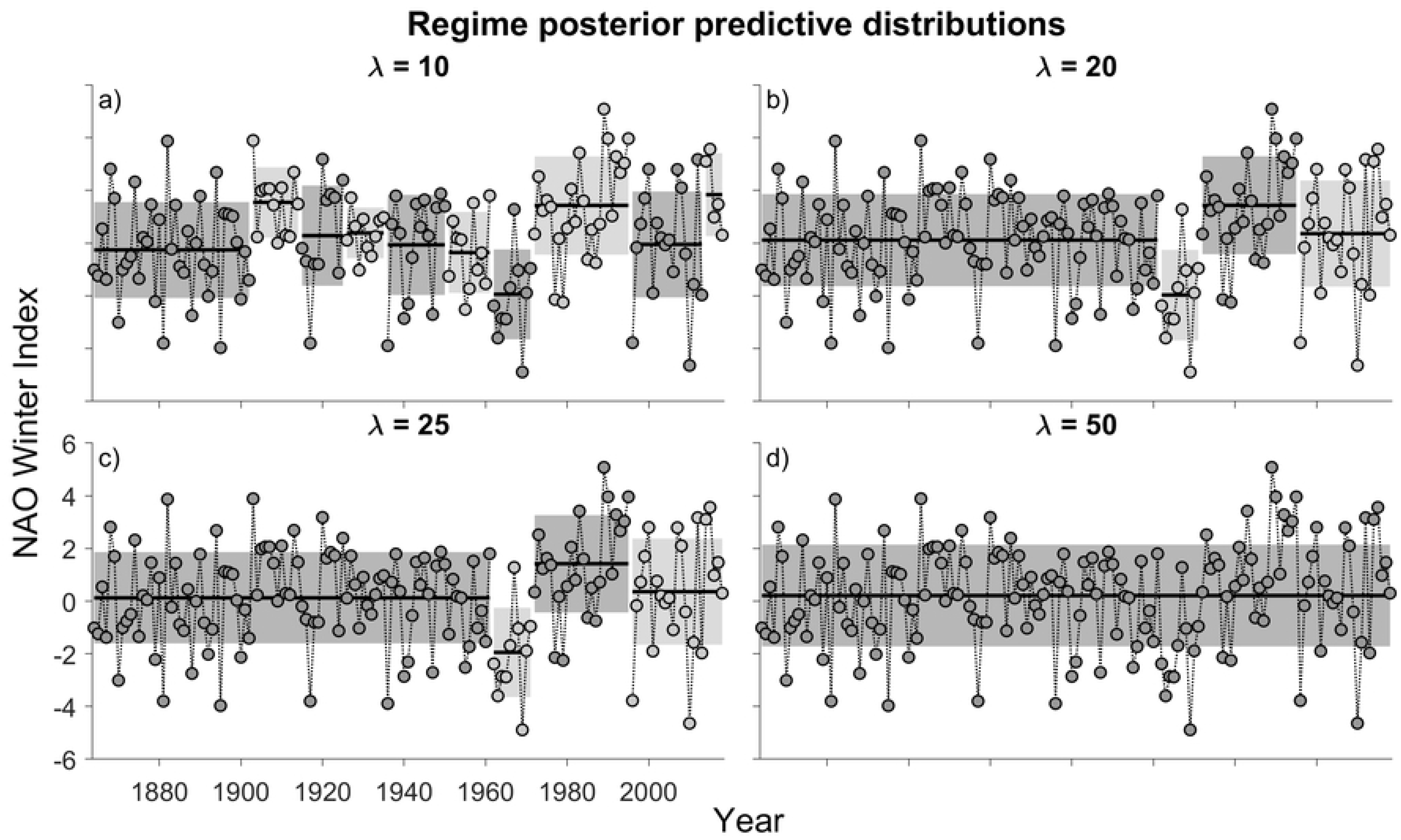
Posterior predictive distributions of time series data on the winter North Atlantic Oscillation (NAO) index. The units are based on the difference of normalized sea level pressure between Lisbon, Portugal and Stykkisholmur/Reykjavík, Iceland. Differently shaded series of data represent different regimes. The shaded area represents the 68% central probability interval (CPI) of the posterior predictive distribution; thus, it includes uncertainty about the mean and the variance. Horizontal lines in each shaded region represents the mean. *M* is the minimum regime length (in years) and λ is the ‘hazard’ rate in the change point prior, i.e., the expectation of the frequency of a regime shift (a λ value of 20 would imply an expectation that regime shifts occur every 20 years).

Taken together, these observations suggest that λ=10 might represent an unduly liberal frequency of break-point changes in the data, resulting in a tendency to ‘over-identify’ regime shifts. To guard against this possibility, we steadily increased λ to as high as 50 years. The model output was identical irrespective of whether λ was set at 20 or 25 years (Figs. 1b,c). Four regimes were detected: 1864-1960 (97 yr); a negative shift from 1961 to 1971 (11 yr); and a positive shift from 1972 to 1995 (24 yr), followed by a negative shift from 1996 to 2018 (23 yr). No regime shifts were detected at λ=50 (Fig 1d), suggesting that this hazard rate is unduly conservative.

### Juvenile Atlantic Cod

At a hazard rate of λ=10, seven regimes of cod catch rate were detected from 1919 to 2014 (Fig 2a). Despite some differences in the timing of the shifts, they, as do all outputs from λ=10 to 50, illustrate patterns of substantive decline. However, there is reason to believe that a value of 10 for the hazard rate yields regime shifts early in the data series that should be interpreted with caution. The first thirty years are characterized by highly variable catch rates. Data from the late 1930s to the mid-1940s seem particularly suspect, given that catch rates between successive years would normally be expected to be positively (not negatively) autocorrelated (the juvenile cod group includes primarily ages 0+ and 1+ yr). The 1939-1945 period coincided with WWII during which fisheries research was greatly curtailed. We note that other researchers (e.g., [16]) excluded this time period from their analyses of the same beach-seine survey data because sampling stations were few in number. Although we do not wish to exclude data, we are disinclined to accept these unusual patterns of data variability from the 1920s to the mid 1940s as reflecting valid regime shifts (recall that the BOCPD algorithm recognizes changes in variability, independent of the mean, as a basis for a regime shift).

**Fig 2.**
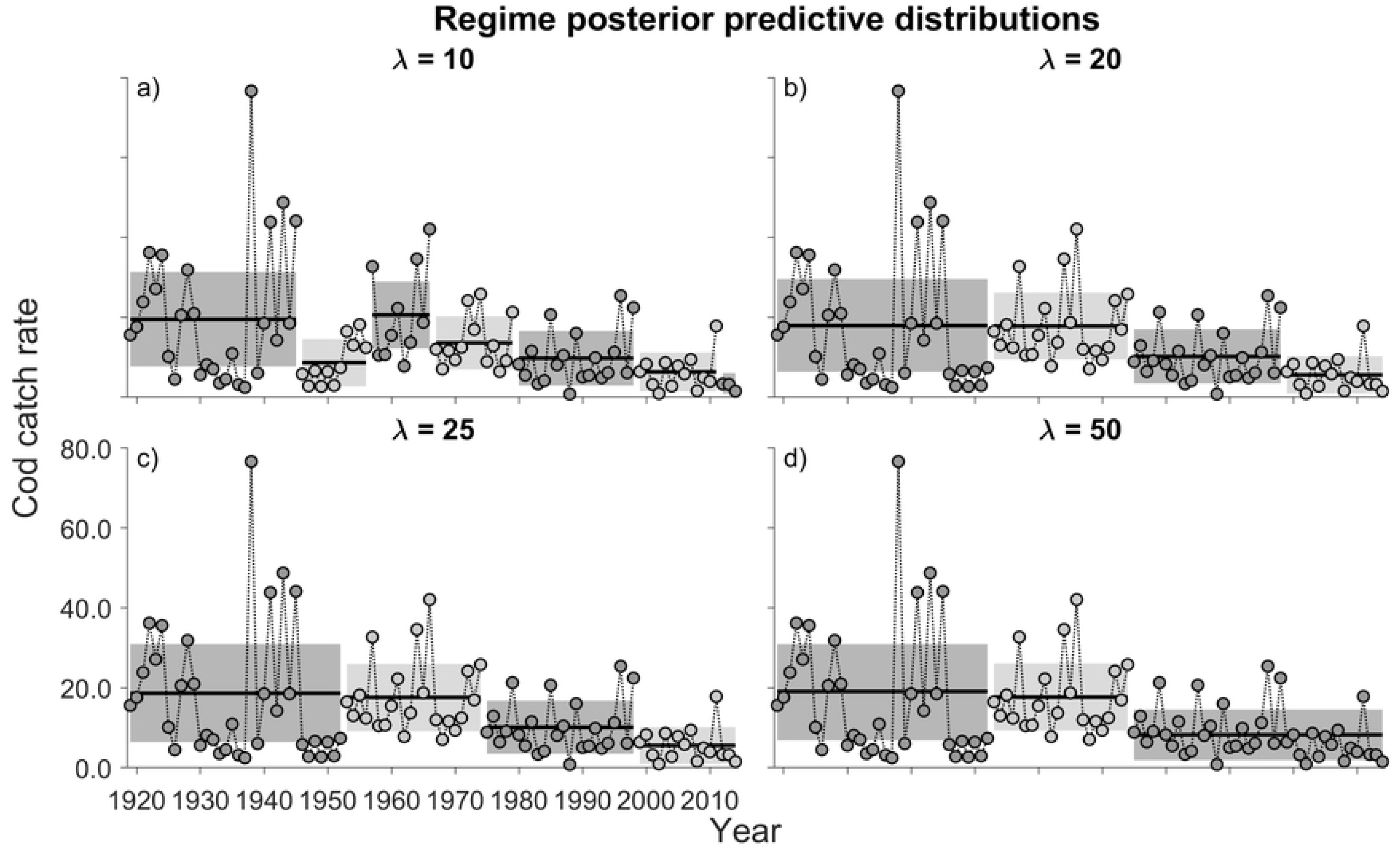
Posterior predictive distribution of juvenile Atlantic cod catch rate (number of juvenile cod per beach-seine haul). See the caption for Fig 1 for explanations of the shaded regions of the data.

As with the NAO index (Figs. 1b,c), λ values of 20 and 25 detected identical regimes (Figs. 2b,c) for the time-series of cod catch rate. Four can be identified. The first two (1919-1952 and 1953-1974) differ only in their variability, the former being more variable than the latter. Catch rates for both regimes were ~18.5 cod per seine haul. However, catch rates during the third (1975-1998) and fourth regimes (1999-2014) averaged ~10 and ~5 cod per haul, respectively. Comparing catch rates in the most recent regime with those in the first two regimes, juvenile cod are estimated to have declined more than 70% since the early 20^th^ Century.

### Zooplankton: Calanus finmarchicus

Unlike the data on the NAO and cod, regime shifts in the abundance of *C. finmarchicus* changed little with changes in λ (Fig 3). At hazard rates of 10 through 50, four regimes were distinguished. Focusing on those at λ=25, the earliest supported the highest average yearly abundance of *C. finmarchicus* in the time series, declining by ~75% in the second regime (1982-1996). The yearly abundance during the third regime (1997-2007) declined further still, before increasing back to the level of the second regime during the fourth regime shift (2008 to 2017).

**Fig 3.**
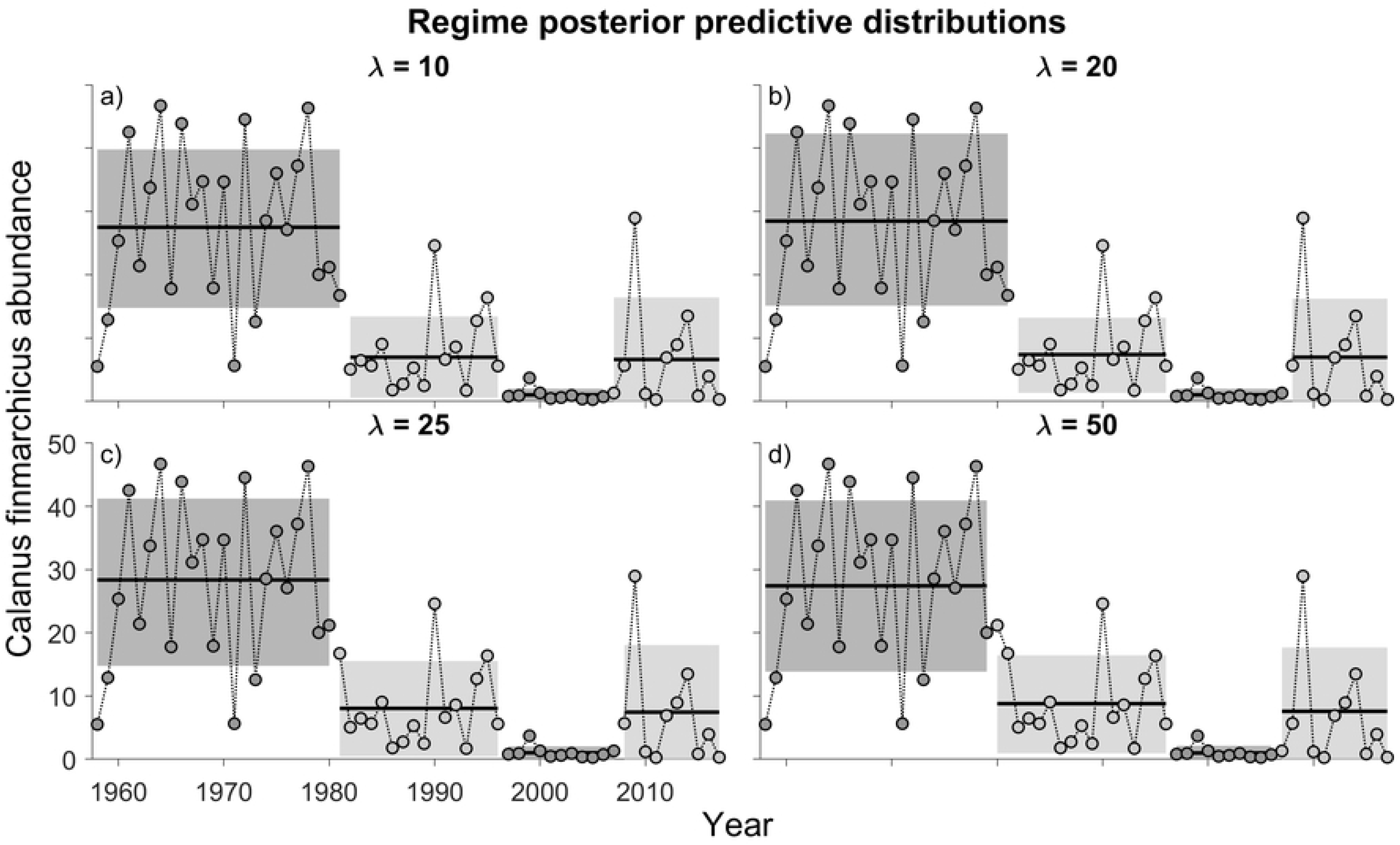
Posterior predictive distributions of data on the abundance of *Calanus finmarchicus* as estimated from the Continuous Plankton Recorder Survey (area C1). Data have been pooled for the months of March through August. Keeping *M* constant at 10 yr, model output is shown for λ values of (a) 10, (b) 20, (c) 25, and (d) 50 years. See the caption for Fig 1 for explanations of the shaded regions of the data.

### Water Temperature

Data on water temperatures are presented as monthly averages (Fig 4 and S1-S3 Figs). At λ values of 20 and 25, the model outputs for the first half of the year were remarkably similar, particularly at the higher hazard rate (λ=25). Two regimes were generally identified with a shift detectable in 1988 for all months from January through June (Fig 4). The mean temperature increase between regimes ranged between 1° and 2°C. As with the first half of the year, two regimes were generally distinguished from July through December, again involving temperature increases of between 1° and 2°C (Fig 4). Interestingly the regime shifts (λ=25) during the latter half of the year tended to occur progressively later than those from January through June: July (1997), August (1994), September (1996), October (1999), November (2005), December (2014). The greatest increase occurred in August, with mean temperatures between the first and second regimes increasing from 16.7° to 18.7°C.

**Fig 4.**
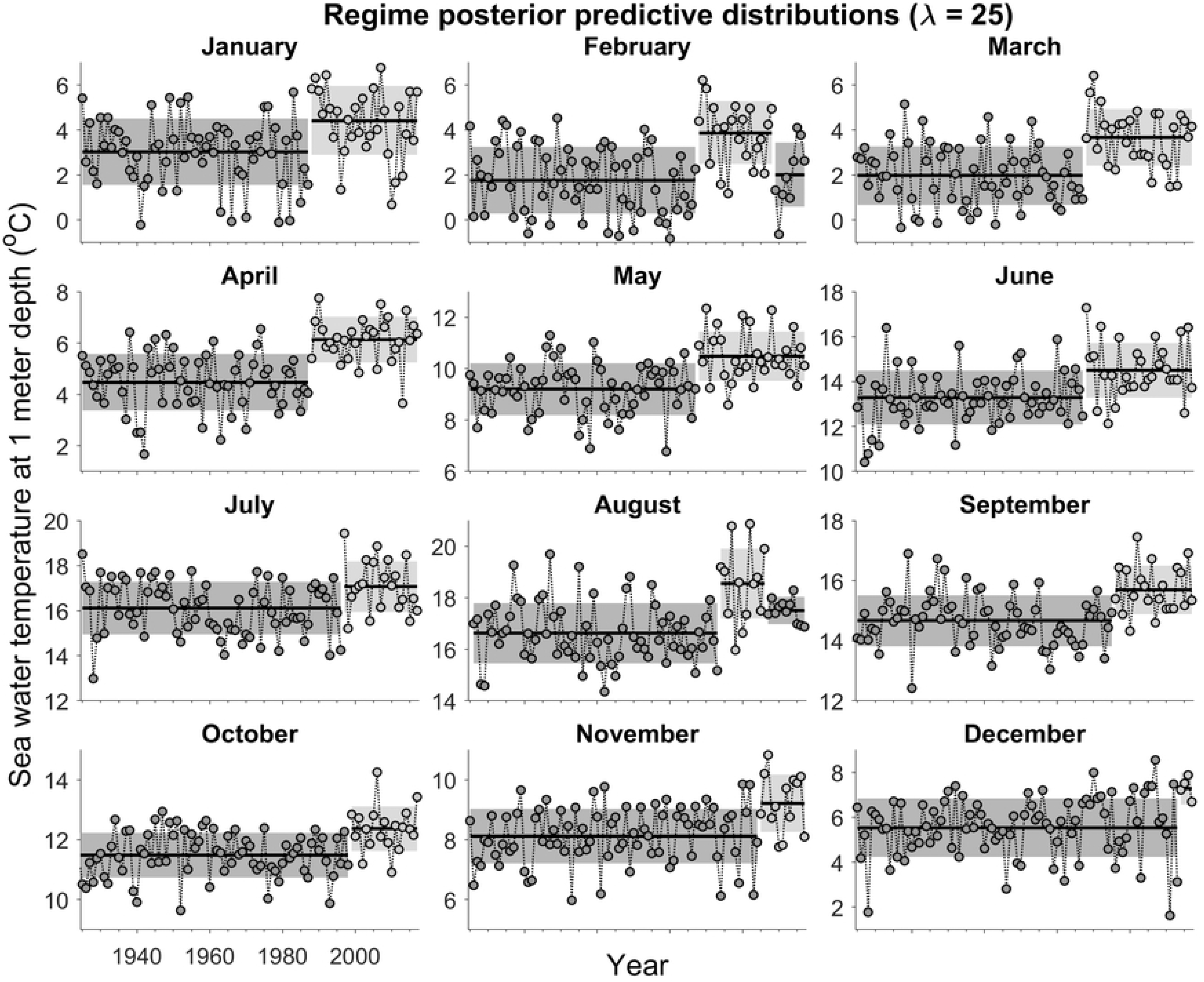
Posterior predictive distributions of data on sea surface water temperatures measured at Flødevigen Research Station for the months of January through December at a hazard rate (λ) of 25 years (*M* = 10 years). See the caption for Fig 1 for explanations of the shaded regions of the data.

### Fishing Mortality and Spawning Stock Biomass

Regime shifts in juvenile cod abundance in Norwegian Skagerrak did not occur in an environment in which cod mortality was affected solely by natural causes. It is well established that fishing can dominate other sources of change in the mean and variance of population size [28]. In the present context, between 1963 and 2017, the mean instantaneous rate of fishing mortality (*F*) on North Sea cod aged 2-4 yr, of which Skagerrak cod are a part, was 0.82 [27]. By contrast, the average annual natural mortality of cod comprising >90% of the catch ranged between 0.2 (cod older than 3 yr) and 0.74 (2-yr-old cod). Thus, since at least the early 1960s, fishing mortality experienced by North Sea cod has always exceeded natural mortality.

Changes in fishing mortality (*F*) and the reproductive component of North Sea cod (the spawning stock biomass, or SSB) can be expressed relative to their limit reference points. When SSB falls below its limit reference point (*B*_*lim*_), the population is considered to have increased risk of impaired reproductive capacity [29]. *F* should not exceed *F*_*lim*_ because such a level of prolonged overfishing is thought to be associated with population dynamics that lead to stock collapse. For North Sea cod, *F*_*lim*_ = 0.54. By comparison, the fishing mortality corresponding to the maximum sustainable yield (*F*_*MSY*_) is 0.31 [27].

The BOCPD algorithm was applied to the fishing mortality data for North Sea cod.

Multiple regime shifts were detected, and these were remarkably similar at different hazard rates (Fig 5). Stock assessment modelling output indicates that fishing mortality on North Sea cod has rarely been less than *F*_*lim*_ (Fig 6), steadily rising from the early 1960s through the late 1990s from 0.9 *F*_*lim*_ in 1963 to a maximum of 2.0 *F*_*lim*_ in 1999 (the year that initiated the second Skagerrak cod regime shift). While *F* steadily increased, stock biomass experienced a decline from a peak SSB of 2.00 *B*_*lim*_ in 1971. By 1999, SSB was less than its limit reference point (0.80 *B*_*lim*_), part of a continuing decline that did not halt until 2006 when it had declined to 0.41 *B*_*lim*_.

**Fig 5.**
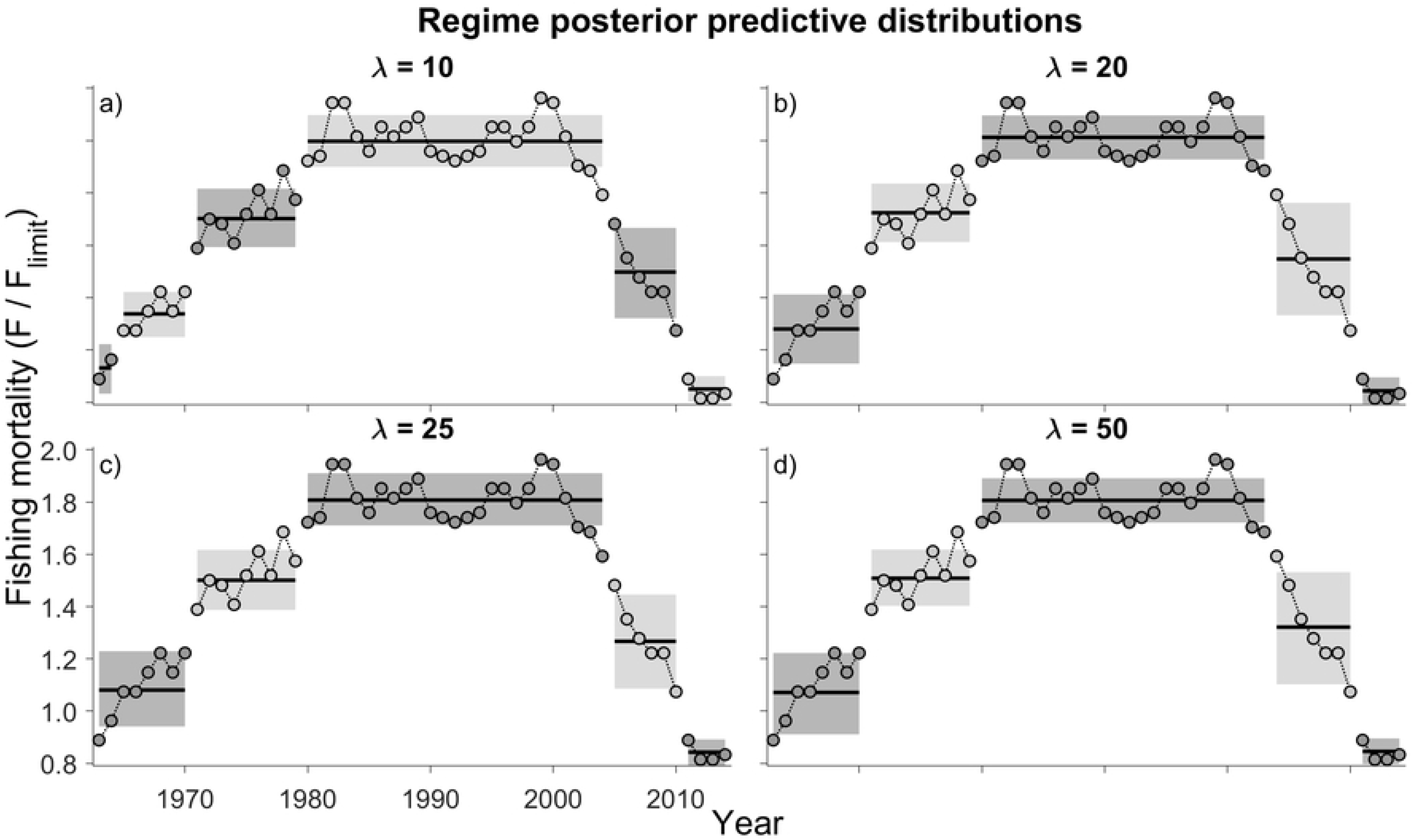
Posterior predictive distributions of estimates of instantaneous rate of fishing mortality (*F*) for cod aged 2-4 yr in the North Sea. Hazard rate (λ) is 10, 20, 25, and 50 years; *M* is held constant at 6 years. See the caption for Fig 1 for explanations of the shaded regions of the data.

**Fig 6.**
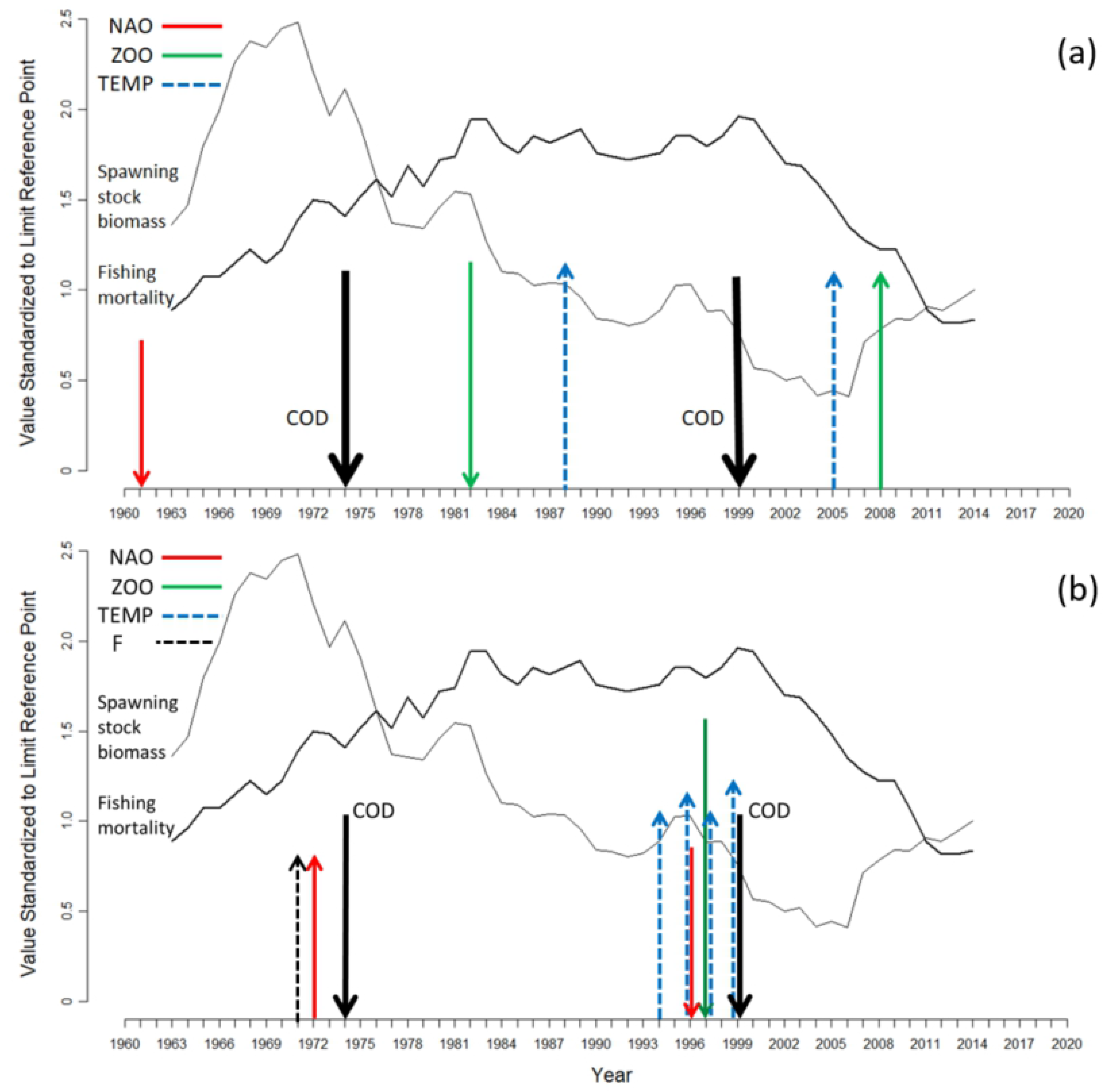
The timing of regime shifts in cod catch rate, NAO, water temperature, and abundance of *Calanus finmarchicus*. Each arrow identifies the beginning of a regime. The direction of the arrow indicates the change in the mean value of the data following each regime shift. The timing of regime shifts is shown in relation to changes in fishing mortality and spawning stock biomass of North Sea cod (each of which is expressed relative to its respective limit reference point; the limit fishing mortality is 0.54, the limit spawning stock biomass is 107,000 tonnes). (a) This panel identifies regime shifts in climate and environmental indices that do not appear to have influenced regime shifts in cod catch rate (i.e., were not followed within 5 years by a cod regime shift). The water temperature regime in 1988 is for January through June; the regime shift in 2005 is for November temperatures. (b) This panel identifies regime shifts in climate and environmental indices that do appear to have influenced regime shifts in cod catch rate (i.e., were followed within 5 years by a cod regime shift). The water temperature regime shifts are for the months of August (1994), September (1996), July (1997), and October (1999).

### Temporal Overlap Between Regime Shifts in Cod Productivity and Putative Drivers

Regime shifts were detected in multiple hypothesized drivers of cod productivity. However, not all of these had obvious, or at least immediate, consequences for cod. That is, regime shifts in NAO, zooplankton abundance, and water temperature were not always followed within 5 years by regime shifts in cod productivity (Fig 6a).

The 1961 NAO regime shift (a decline from 0.1 to −2.0) had no discernable impact, although the cod regime shift that began in 1974 was preceded by regime-shift increases in both NAO (−2.0 to 1.5) and fishing mortality (1.10 to 1.52 *F*_*lim*_). The initial ~75% reduction in copepod abundance in 1981 followed, rather than preceded, the 1974 shift in cod by 6-7 years (Fig 6a). Although the further reduction (~90% decline) in *C. finmarchicus* in 1997 preceded the 1999 regime-shift decline of cod (Fig 6b), cod did not respond when zooplankton increased in 2008 back to its level in the 1981-1996 period. The nearly 50% reduction in average cod abundance that began in 1974 was not preceded by a regime shift in water temperature in any month (Fig 6a). However, the 1999 cod regime shift either coincided with (October) or was preceded 2-5 years earlier (July-September) by regime-shift increases in water temperature of 1° to 2°C (Fig 6b), reaching a maximum monthly mean of almost 19° C in August.

## Discussion

The present study identifies regime shifts in a long-term metric of biological productivity in relation to regime shifts in potential causal drivers, using a Bayesian online change point detection (BOCPD) algorithm. Juvenile Atlantic cod in Norwegian coastal Skagerrak experienced two stepwise reductions in mean abundance, beginning in 1975 and 1999, ultimately attaining a level ~30% of what it was in the first half of the 20^th^ Century. Regime shifts in indices for the NAO and zooplankton abundance since 1960 were directionally the same, the opposite, or independent of directional shifts in cod catch rates. The degree of temporal overlap between regimes shifts in cod abundance and water temperature depended on season. Single regime shifts of increased mean temperature (by 1 to 2°C) during the winter and spring months either preceded the 1999 cod regime shift by more than a decade or occurred several years thereafter. However, summer-autumn temperature regime shifts (1 to 2°C increase) were either concomitant with, or occurred slightly in advance of, the cod regime shift (decrease) in 1999. The earlier cod regime shift (1975) was not associated with a regime shift in water temperature for any month of the year.

The relative importance of hypothesized drivers can be ascertained by the temporal proximity of their regime shifts with regime shifts in cod. It is clear that shifts in some factors, such as the NAO index and abundance of *C. finmarchicus*, do not always have obvious biological consequences for Atlantic cod (see also [30]). Against the background of temporal changes in fishing mortality and spawning stock biomass of North Sea cod, the earliest regime shifts in NAO (1961; a decline from 0.2 to −2.0) and *C. finmarchicus* (1982; a ~75% decline) were not linked with regime shifts in cod (Fig 3a). The same was true of the 1-2° C regime-shift increase in water temperature during the winter-spring months in 1988.

One interpretation of this lack of influence is that drivers of cod productivity are less likely to manifest biological change when (i) they act singly, (ii) human-induced mortality is relatively low, and (iii) cod population size is relatively high. When the NAO shifted in 1961, fishing mortality on North Sea cod was less than *F_lim_* and SSB was 1.36 *B*_*lim*_; when zooplankton abundance shifted downwards in 1982, fishing mortality was increasing but population biomass remained high (1.53 *B*_*lim*_). Regarding the winter-spring increase during the regime shift in water temperature that began in 1988, it is notable that, despite these increases, temperatures remained well within the range thought to be optimal for cod eggs and larvae [31, 32]. Indeed, these January-June regime shifts might have had a beneficial influence on cod productivity, mitigating to some extent any steadily increasing effects of persistently high fishing mortality and declining SSB.

Our analyses reveal two occasions when regime shifts in potential drivers of cod productivity preceded regime shifts in cod catch rate by a time period sufficiently brief (≤ 5 yr) that they could plausibly have influenced the subsequent abundance of cod aged 0+ to 2+ years (Fig 6). Even though a decline in NAO in 1961 had no discernable effect on cod, the increase in 1972 might well have, insofar as the first cod regime shift followed two years later. This supposition is supported by studies (e.g., [19]) that have concluded that an increased NAO index has a negative influence on cod productivity in the northeast Atlantic, although this association may be weakening [20]. One possible reason for why the NAO apparently affected cod (beginning in 1974) is that the magnitude of the NAO regime shift (−2.0 to 1.5) was the greatest of the three shifts that occurred between 1864 and 2018. It is also notable, however, that the 1974 cod regime shift occurred during a period of steadily increasing and unsustainably high fishing mortality (1.5 *F*_*lim*_), potentially affecting the ability of cod to resist environmental changes caused by the NAO, changes to which an unfished population might have been resilient. This underscores the challenge in disentangling the effects of fishing and climate-related indices on biological productivity [33].

The cod regime shift that began in 1999 was preceded by a ‘perfect storm’ of multiple concomitant changes in the environment. Summer-autumn temperatures jumped 1-2° C; the NAO index declined from 1.5 to 0.5; *C. finmarchicus* had plunged to its lowest level in the time series; fishing mortality was at its highest level since 1963 (2.0 *F_lim_*); and spawning stock biomass was at its lowest level in the time series (0.8 *B_lim_*) en route to a minimum of 0.41 *B_lim_* in 2006. The effects of the NAO, *C. finmarchicus*, and temperature on cod productivity were undoubtedly accentuated by the directionality of their regime shifts in 1999. As the NAO index declines, so does primary and secondary productivity in Skagerrak [34], and increased water temperatures are associated with increasingly unfavourable conditions for *C. finmarchicus* [16].

Among the putative drivers of cod productivity, the 1996 regime shift in NAO may have been the most benign, given that (i) a reduction in the index did not have its anticipated positive effect on cod and that (ii) the index had returned to levels characteristic of the 1868 to 1960 period. The very considerable reduction in *C. finmarchicus* (beginning in 1997; Fig 3) was likely a much more prominent factor, given the exceedingly low levels to which this key prey species of juvenile cod had declined.

There is, however, reason to believe that the summer-autumn increase in water temperature was of considerably greater importance than either the NAO or zooplankton abundance. Successive regime shifts from 1994 to 1999 during July through October raised temperatures to their highest recorded levels in coastal Skagerrak since 1925, when the time series began. In some years, mean August temperatures exceeded 20°C, approaching the critical thermal maximum for Atlantic cod [32,35].

We hypothesize that cod did not respond positively to the presumed increase in food supply in 2008 because of the physiological stress associated with increased summer-autumn water temperatures. Based on tagging studies at sea of almost 400 cod from 8 northeast Atlantic cod stocks, Righton et al. [33] found that although the total thermal niche of adult cod ranged between −1.5 and 19.0° C, the temperature range was considerably narrower during the spawning period when larval and juvenile cod are developing (1 to 8° C). Nissling [31] reported that survival of larval cod in the laboratory declined considerably when water temperatures exceeded 10° C.

One fundamentally important element to consider when evaluating the consequences of climate-related and environmental regime shifts on population productivity is the size of the population relative to a metric of long-term sustainability, such as carrying capacity or population size in an unfished state. This is because small populations are more vulnerable to environmental stochasticity than comparatively large populations [36]. This link between population size and susceptibility to environmental change has been repeatedly considered when assessing the recovery capacity of depleted cod populations [37–38]. But it has also been made with respect to potential drivers of cod regime shifts. Based on an analysis of cod populations on the European Shelf south of 62°N, including North Sea cod, Brander [39] concluded that environmental variability, as represented by the NAO index, only affects cod when the spawning stock biomass is low. Brander’s [39] argument is both theoretically compelling and empirically supported by the present study.

There are several attributes to the methodology we have applied here. Firstly, the same algorithm is used to identify regime shifts in a metric of biological productivity and putative causal drivers of that metric. Secondly, our approach greatly reduces the subjectivity inherent in deciding the magnitude of data change which constitutes a regime shift (the ‘effect’ size) and when it is that a regime shift occurs; we did not presume the existence of a regime shift in any given year for any given variable. A third improvement is that the BOCPD algorithm accounts for changes in the variance in the data, not simply the mean.

One limitation in our interpretation of the relative importance of fishing and the environment on regime shifts in cod productivity is our use of estimates of fishing mortality and spawning stock biomass for North Sea cod as metrics of fishing pressure and population size for Skagerrak cod. But if we were to account for fishing mortality in our analyses, we needed to avail ourselves of the best available data in this regard, and these data were available for North Sea cod. There are empirically defensible reasons for our application of North Sea cod estimates of *F* and SSB to Skagerrak cod. Firstly, North Sea cod genotypes exist along the Norwegian Skagerrak coast [40]. Secondly, Skagerrak has long been considered part of the North Sea cod stock unit [27]. Thirdly, limited estimates of fishing mortality available for Skagerrak cod confirm that fishing mortality can be exceedingly high. Kleiven et al. [41] reported that recreational and commercial fisheries for cod in Skagerrak fjords resulted in a mortality rate of 55.6% for the years 2005 to 2013, equivalent to *F*=0.81. For comparison, the average *F* for North Sea cod over the same time period was 0.61 [27], suggesting that the fishing mortality experienced by North Sea cod may be comparable to, and possibly less than, that experienced by Norwegian Skagerrak coastal cod in some years.

The concept of regime shifts permeates the marine ecological and fisheries literature. Definitions vary considerably. The ecological literature tends to interpret regime shifts as community-level changes between alternative stable states with the implication that such shifts are difficult to reverse [5, 7]. In contrast, regime-shift analyses of meteorological factors tend not to focus on alternative stable states, being much more accepting of regime-shift ‘reversibility’ (e.g., [12, 20]). The fisheries literature is perhaps intermediate with respect to the question of regime-shift reversibility. Some work draws attention to long-term, slow-to-reverse discontinuities in ecosystem properties [4], whereas neither reversibility nor regime-shift time period have been integral to a lack of temporal stationarity in fish-stock productivity (e.g., [11, 14]).

Our analyses emphasize the utility in examining multiple regime shifts when trying to understand the causal mechanisms responsible for regime shifts in metrics of biological productivity. Doing so allows one to formulate hypotheses and to draw conclusions concerning the conditional probabilities that an environmentally related regime shift will affect biological productivity. One hypothesis that emerges here is that the strength of the effect of an environmental or climate-related regime shift is accentuated when it coincides with other regime shifts. A second hypothesis, underscoring the findings of previous work [38–39], is that climate-related regime shifts are more likely to affect populations when they are relatively small. The present study affirms the dominant role that fishing has on the probability that populations will respond to regime shifts in environmental variables, underscoring the fundamental necessity of accounting for fishing mortality in any analysis of regime shifts in commercially exploited marine fishes [4, 18, 33].

For our case study of Norwegian Skagerrak cod, our work suggests that steadily increasing fishing mortality from commercial and recreational fisheries has increasingly sensitized the cod to regime shifts in NAO, zooplankton abundance, and water temperature. Fishing mortality remains unsustainably high in the region [41]. This, coupled with small population size and increased summer and autumn water temperatures that broach the thermal limit for the species, are likely major factors limiting the recovery capacity for cod in southern coastal Norway.

## Acknowledgements

The authors are very grateful to the Sir Alister Hardy Foundation for Ocean Science (SAHFOS, Marine Biological Association, Plymouth, UK, for their permission to analyze data from the Continuous Plankton Recorder Survey. We thank, in particular, Martin Edwards and David Johns of SAHFOS.

## Supporting information

S1 Fig. Posterior predictive distributions of data on sea surface water temperatures measured at Flødevigen Research Station for the months of January through December at a hazard rate (λ) of 10 years (*M* = 10 years). The shaded area represents the 68% central probability interval (CPI) of the posterior predictive distribution; thus, it includes uncertainty about the mean and the variance. Horizontal lines in each shaded region represents the mean.

S2 Fig. Posterior predictive distributions of data on sea surface water temperatures measured at Flødevigen Research Station for the months of January through December at a hazard rate (λ) of 20 years (*M* = 10 years). See the caption for Fig 1 or S1 Fig for explanations of the shaded regions of the data.

S3 Fig. Posterior predictive distributions of data on sea surface water temperatures measured at Flødevigen Research Station for the months of January through December at a hazard rate (λ) of 50 years (*M* = 10 years). See the caption for Fig 1 or S1 Fig for explanations of the shaded regions of the data.

